# Co-targeting B-RAF and PTEN enables sensory axons to regenerate across and beyond the spinal cord injury

**DOI:** 10.1101/2022.03.07.483366

**Authors:** Harun N. Noristani, Hyukmin Kim, Shuhuan Pang, Jian Zhong, Young-Jin Son

**Affiliations:** Shriners Hospitals Pediatric Research Center and Center for Neural Repair, Lewis Katz School of Medicine, Temple University, Philadelphia, PA 19140, USA; Burke Medical Research Institute, Weill Cornell Medical College of Cornell University, White Plains, NY 10605, USA

**Keywords:** Spinal cord injury, primary afferents, glial scar, DRG, dorsal column axons, sensory axon regeneration, conditioning lesion

## Abstract

Primary sensory axons in adult mammals fail to regenerate after spinal cord injury (SCI), in part due to insufficient intrinsic growth potential. Robustly boosting their growth potential continues to be a challenge. Previously, we showed that constitutive activation of B-RAF (rapidly accelerated fibrosarcoma kinase) markedly promotes axon regeneration after dorsal root and optic nerve injuries. The regrowth is further augmented by supplemental deletion of PTEN (phosphatase and tensin homolog). Here, we examined whether concurrent B-RAF activation and PTEN deletion promotes dorsal column axon regeneration after SCI. Remarkably, genetically targeting B-RAF and PTEN selectively in DRG neurons of adult mice enables many DC axons to enter, cross, and grow beyond the lesion site after SCI; some axons reach ~2 mm rostral to the lesion by 3 weeks post-injury. Co-targeting B-RAF and PTEN promotes more robust DC regeneration than a pre-conditioning lesion, which additively enhances the regeneration triggered by B-RAF/PTEN. We also found that post-injury targeting of B-RAF and PTEN enhances DC axon regeneration. These results demonstrate that co-targeting B-RAF and PTEN effectively enhances the intrinsic growth potential of DC axons after SCI and therefore may help to develop a novel strategy to promote robust long-distance regeneration of primary sensory axons.

## INTRODUCTION

Adult mammalian central nervous system (CNS) neurons fail to regenerate their axons after spinal cord injury (SCI). This regeneration failure is due to extrinsic inhibitory cues and glial scar formation that hinder axonal re-growth by imposing physical and chemical barriers [for reviews see (Silver and Miller, 2004, Geoffroy and Zheng, 2014)]. The lack of a sufficiently robust intrinsic regenerative response also substantially contributes to the inability of adult CNS neurons to regenerate their severed axons across and beyond the lesion site [for reviews see (Mar et al., 2014, He and Jin, 2016, Curcio and Bradke, 2018)].

The pseudounipolar sensory neurons within the dorsal root ganglia (DRG) project a peripheral branch that innervates muscle, skin, and joints and a central branch [also referred to as dorsal column (DC) axons] that transmits information to the spinal cord and brainstem. In adults, a lesion to the peripheral branch prior to a lesion to the central branch, known as a pre-conditioning lesion, initiates an intrinsic regenerative response that enhances limited regeneration of DC axons (McQuarrie and Grafstein, 1973, Richardson and Verge, 1986, Neumann and Woolf, 1999, Richardson and Issa, 1984).

Studies over the past decade have identified several molecules that regulate intrinsic regeneration, including: phosphatase and tensin homolog (PTEN) (Park et al., 2008, Liu et al., 2010), rapidly accelerated fibrosarcoma kinase (B-RAF) (O’Donovan et al., 2014), suppressor of cytokine signaling 3 (SOCS3) (Smith et al., 2009), Krüppel-like factors (KLFs) (Blackmore et al., 2012, Moore et al., 2009), c-myc (Belin et al., 2015), SRY-Box Transcription Factor 11 (*Sox11*) (Wang et al., 2015, Norsworthy et al., 2017), and Lin28 (Wang et al., 2018, Nathan et al., 2020). However, the limited efficacy with which these molecules promote DC axon regeneration following SCI emphasizes the urgent need to identify additional molecules that can powerfully stimulate robust, lengthy axon re-growth.

B-RAF is critical for neurotrophin-induced sensory axon outgrowth (Markus et al., 2002, Zhong et al., 2007). We demonstrated previously that constitutive, selective activation of B-RAF in adult DRG neurons enables their axons to regenerate into the spinal cord in adult mice after dorsal root injury (O’Donovan et al., 2014). B-RAF activation also enhances regeneration of the optic nerves. Importantly, concomitant B-RAF activation and PTEN deletion synergistically increases regeneration of both optic nerve (O’Donovan et al., 2014) and dorsal root (Kim, Noristani et al., in preparation). No studies have yet examined the effects of concurrently targeting B-RAF and PTEN on DC axon regeneration after SCI. Here we demonstrate that simultaneous B-RAF activation and PTEN deletion enables DC axons to regenerate up to 2000 μm rostral to the lesion site by 3 weeks post SCI. The extent of this regrowth is significantly greater than that enabled by a pre-conditioning lesion, which, however, additively enhances the regeneration stimulated by B-RAF/PTEN. We also report that concomitant B-RAF activation and PTEN deletion immediately after SCI promotes DC axon regeneration. These results identify concurrent B-RAF activation and PTEN deletion as a promising strategy to heighten intrinsic regeneration of adult sensory neurons and to promote axon regeneration after SCI.

## RESULTS

### Concomitant B-RAF activation and PTEN deletion promotes dorsal column axon regeneration across the spinal cord lesion

To examine whether simultaneous B-RAF activation and PTEN deletion enhances DC axon regeneration after SCI, we used inducible transgenic mice expressing constitutively active B-RAF with PTEN deletion selectively in dorsal root ganglia (DRG) neurons: *LSL-kaBRAF:PTEN^f/f^:Brn3a-CreER^T2^:R26-TdT* (hereafter as kaBRAF/PTEN iTg). We used 6-weeks-old kaBRAF/PTEN iTg and age-matched *Brn3a-CreER^T2^*-negative littermates as controls (WT). To induce kaBRAF expression and PTEN deletion, we administered tamoxifen for 5 consecutive days, 2 weeks prior to SCI (Figure 1A). We visualized ascending DC axons by microinjecting the recombinant self-complementary Adeno-Associated Virus serotype 2 expressing enhanced green fluorescent protein (scAAV2-eGFP) into the lumbar level (L) 4 and L5 DRGs. Recently, we have shown that AAV2-eGFP injections in the DRG predominantly label proprioceptive and mechanoreceptive axons (Zhai et al., 2021). Using GFAP immunostaining, we first assessed the lesion area in all sagittal sections with DC axons. The lesion area ranged between 0.093 – 0.192 mm^2^ in WT and 0.028 – 0.172 mm^2^ in kaBRAF/PTEN iTg mice (Figure 1B), suggesting that our SCI model produced relatively wide lesions. Mice with unusually enlarged or narrow lesion areas were excluded from the analysis. The lesion width at the injury epicenter ranged between 145 – 230 μm in WT and 100 – 220 μm in kaBRAF/PTEN iTg mice. Quantification of lesion area showed no significant differences between WT and kaBRAF/PTEN iTg groups (Figure 1B). We then determined the total number of regenerating DC axons at specific distances rostral to the lesion epicenter using a consecutive series of GFP and GFAP overlay images (Figure 1C&D, Supplementary Figure 1).

**Figure 1.**
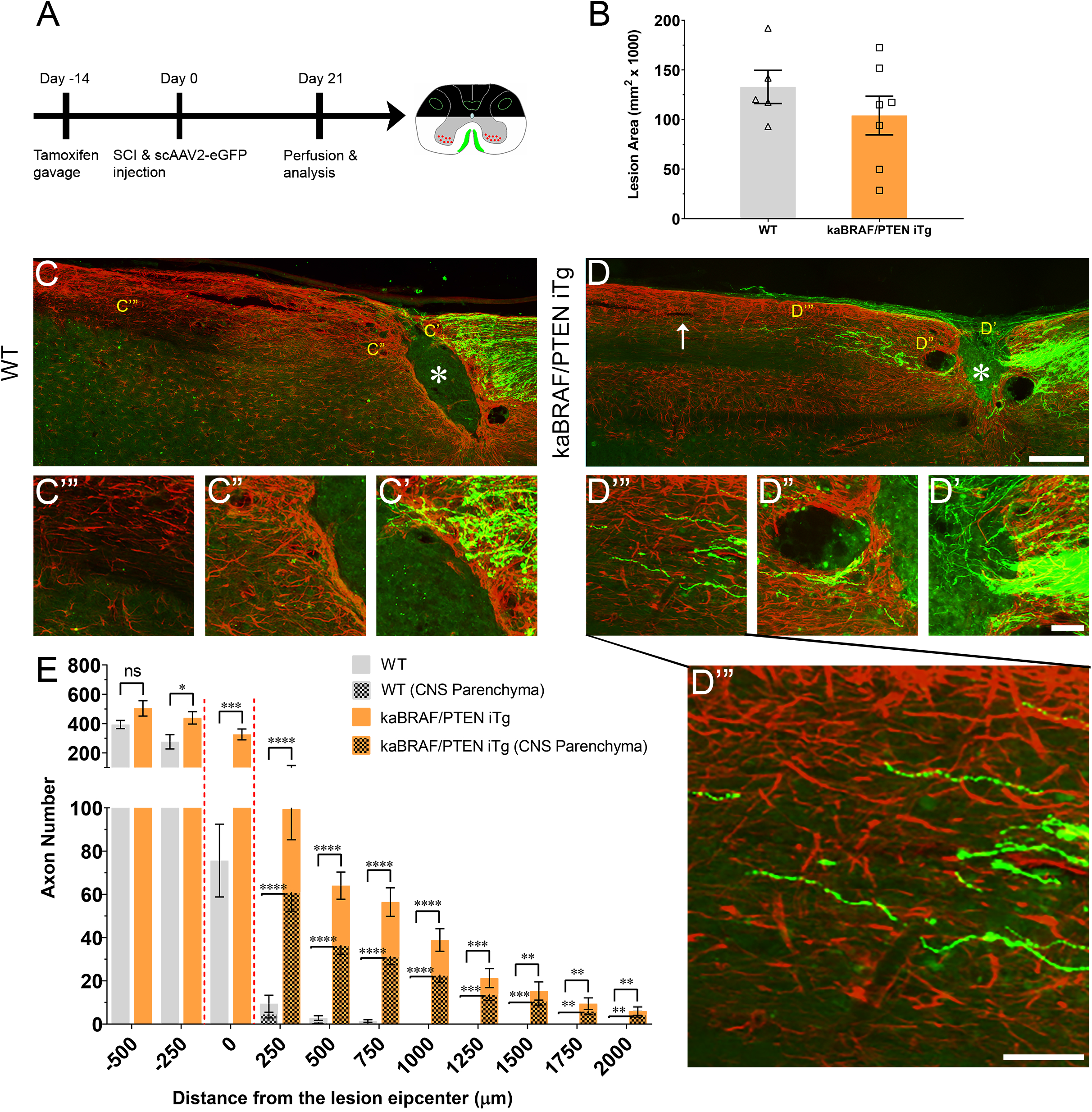
Concomitant B-RAF activation and PTEN deletion promotes DC axon regeneration. (A) Schematic diagram showing the experimental timeline. Two weeks after the first tamoxifen gavage WT and kaBRAF/PTEN iTg mice underwent dorsal SCI and scAAV2-eGFP injection into the right lumbar (L) 4 and L5 DRGs to label regenerating DC axons. (B) Quantitative analysis of the lesion area after dorsal SCI in kaBRAF/PTEN iTg versus WT mice. (C–C’”) Representative images of a WT mouse spinal cord showing most DC axons stopping at the caudal astroglial border of the injury site and very few growing along the astrocytic bridge that also terminates ~200 μm from the lesion epicenter. (D–D’”) Representative images of a kaBRAF/PTEN iTg mouse spinal cord showing increased DC axon regeneration. Many regenerating DC axons penetrate the lesion epicenter; several extend directly across the rostral astroglial border of the injury site. (E) Quantitative analysis of total numbers of regenerated DC axons in kaBRAF/PTEN iTg versus WT mice. DC axon regeneration is significantly greater in kaBRAF/PTEN iTg mice at multiple distances measured from the lesion epicenter: −250μm: *P=0.0133, df=10; 0μm: ***P=0.0009, df=10; 250μm: ****P<0.0001, df=10; 500μm: ****P<0.0001, df=10; 750μm: ****P<0.0001, df=10; 1000μm: ****P<0.0001, df=10; 1250μm: ***P=0.0003, df=10; 1500μm: **P=0.0029, df=10; 1750μm: **P=0.0037, df=10; 2000μm: **P=0.0081, df=10; two-way ANOVA with repeated measures. ~75% of regenerating DC axons in kaBRAF/PTEN iTg mice enter the CNS parenchyma (the area with GFAP-positive astrocytes); the remaining 25% grow along the spinal cord surface. (E, patterned) Quantitative comparison of DC axons regenerated into CNS parenchyma in kaBRAF/PTEN iTg versus WT mice. Regenerating DC axons within CNS parenchyma are significantly increased at several distances rostral to the SCI in kaBRAF/PTEN iTg mice. 250μm: ****P<0.0001, df=10; 500μm: ****P<0.0001, df=10; 750μm: ****P<0.0001, df=10; 1000μm: ****P<0.0001, df=10; 1250μm: ***P=0.0004, df=10; 1500μm: ***P=0.0003, df=10; 1750μm: **P=0.0034, df=10; 2000μm: **P=0.0053, df=10; two-way ANOVA with repeated measures. n=5 (WT), and 7 (kaBRAF/PTEN iTg). Scale bars, 250μm (C and D), 50μm (C’–C’”, and D’–D’”).

In WT mice most DC axons stopped at the astroglial border caudal to the injury site with very few axons growing over the astrocytic bridge and then abruptly stopping within 200 μm of the lesion epicenter (Figure 1C). DC axons in WT mice displayed characteristic dieback from the lesion epicenter, and individual axons had several retraction bulbs that appeared as bulbous swellings in their axonal tips (Figure 1C–C’). In all WT mice examined, DC axons failed to reach the rostral border of the lesion (Figure 1C”) or extend rostrally across the astroglial border (Figure 1C’”). Remarkably, in kaBRAF/PTEN iTg mice numerous DC axons crossed the caudal astroglial border and penetrated the lesion epicenter (Figure 1D). In most kaBRAF/PTEN iTg mice, regenerating DC axons that entered the lesion epicenter appeared preferentially to grow on the surface of the spinal cord (Figure 2D). Regenerating DC axons extending along the spinal cord surface frequently penetrated the CNS parenchyma rostral to the lesion site (Figure 2D’”). In a few kaBRAF/PTEN iTg mice, dramatic rostral growth was also found across the astroglial border (Figure 1D”–D’”). DC axons in the kaBRAF/PTEN iTg group exhibited a characteristic regenerative morphology with tortuous trajectories (Figure 1D). Most DC axonal tips in kaBRAF/PTEN iTg mice were thin and continuous with the axonal shaft resembling growth cones (Figure 1D’–D’”). Quantification revealed a significant increase in regenerating DC axons >2000 μm rostral to the lesion site in kaBRAF/PTEN iTg mice compared to the WT group (Figure 1E). Using GFAP immunostaining, we next specifically examined regenerating DC axons within the CNS parenchyma. Around two-thirds of regenerating DC axons in kaBRAF/PTEN iTg mice appeared within the CNS parenchyma i.e., the area with GFAP-positive astrocytes, while the remaining regenerating DC axons projected along the spinal cord surface. Quantification of regenerating DC axons within the CNS parenchyma confirmed a significant increase >2000 μm across the lesion epicenter in kaBRAF/PTEN iTg mice compared to the WT group (Figure 1E). Therefore, concomitant B-RAF activation and PTEN deletion powerfully promotes regeneration of DC axons, enabling them to extend up to ~2 mm beyond the lesion within 3 weeks after SCI.

**Figure 2.**
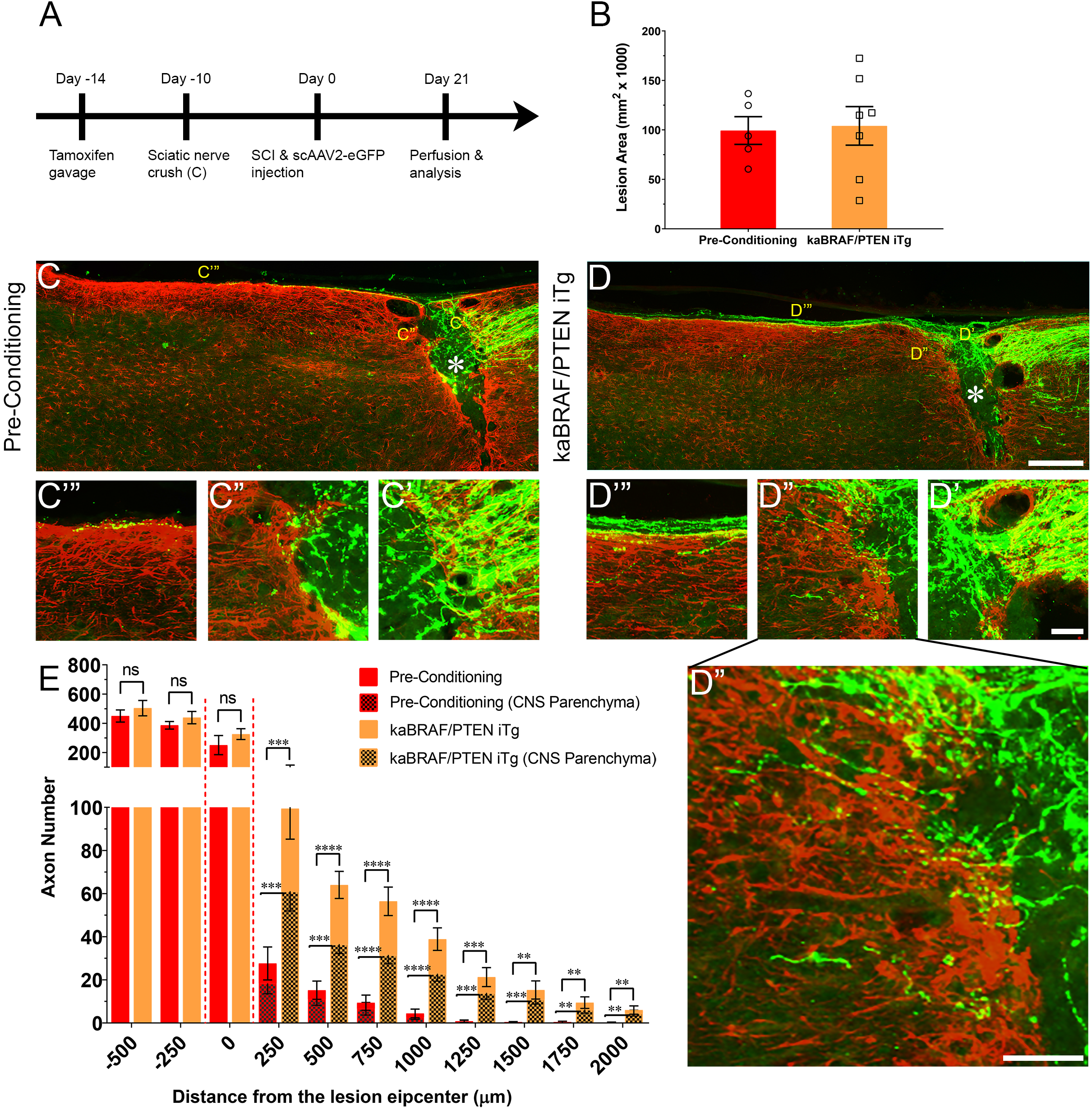
Concomitant B-RAF activation and PTEN deletion enhances DC axon regeneration more effectively than a pre-conditioning lesion. (A) Schematic diagram showing the experimental timeline. Four days after the first tamoxifen gavage one group of WT mice underwent right sciatic nerve crush (pre-conditioning lesion). WT mice with a pre-conditioning lesion and kaBRAF/PTEN iTg mice without pre-conditioning lesion underwent dorsal SCI and scAAV2-eGFP injection into the right L4 and L5 DRGs to label regenerating DC axons two weeks after the first tamoxifen gavage. (B) Quantitative analysis of the SCI lesion area in kaBRAF/PTEN iTg group versus pre-conditioning group. (C–C’”) Representative images of a pre-conditioning mouse spinal cord showing regenerating DC axons reaching the lesion epicenter with a few terminating just rostral to the lesion site. (D–D’”) Representative images of a kaBRAF/PTEN iTg mouse spinal cord showing robust DC axon regeneration. Many regenerating DC axons penetrate the lesion epicenter and extend long distances rostral to the lesion site. (E) Quantitative comparison of regenerated DC axons in kaBRAF/PTEN iTg mice versus WT pre-conditioning lesion group. Significantly greater DC axon regeneration is evident at multiple distances rostral to the lesion in kaBRAF/PTEN iTg mice. 250μm: ***P=0.0004, df=10; 500μm: ****P<0.0001, df=10; 750μm: ****P<0.0001, df=10; 1000μm: ****P<0.0001, df=10; 1250μm: ***P=0.0005, df=10; 1500μm: **P=0.0035, df=10; 1750μm: **P=0.005, df=10; 2000μm: **P=0.0100, df=10; two-way ANOVA with repeated measures. (E, patterned) Quantitative analysis of regenerated DC axons in CNS parenchyma rostral to the injury site in kaBRAF/PTEN iTg mice versus WT pre-conditioning lesion group. Significantly increased DC axon regeneration is evident at multiple distances up to 2000 μm rostral to the lesion in kaBRAF/PTEN iTg mice. 250μm: ***P=0.0008, df=10; 500μm: ***P=0.0001, df=10; 750μm: ****P<0.0001, df=10; 1000μm: ****P<0.0001, df=10; 1250μm: ***P=0.0006, df=10; 1500μm: ***P=0.0004, df=10; 1750μm: **P=0.0054, df=10; 2000μm: **P=0.0071, df=10; two-way ANOVA with repeated measures. n=5 (pre-conditioning), and 7 (kaBRAF/PTEN iTg). (F–G). Scale bars, 250μm (C and D), 50μm (C’–C’”, and D’–D’”), ns, not significant.

### Concomitant B-RAF activation and PTEN deletion enhances DC axon regeneration more effectively than a pre-conditioning lesion

It is firmly established that sciatic nerve injury enhances the regenerative capacity of adult DRG neurons following a subsequent injury of DC axons. This injury is known as a ‘pre-conditioning’ lesion (Richardson and Issa, 1984, Richardson and Verge, 1986). To compare the extent of DC axon regeneration between kaBRAF/PTEN iTg mice and mice with a pre-conditioning lesion, we crushed the right sciatic nerve of another group of WT mice 10 days before SCI (Figure 2A). The lesion area in pre-conditioning mice ranged between 0.06 – 0.137 mm^2^ and was not significantly different from that in kaBRAF/PTEN iTg mice (Figure 2B). Like kaBRAF/PTEN iTg mice, all mice examined in the pre-conditioning group showed numerous DC axons that crossed the caudal astroglial border and entered the lesion epicenter. However, they rarely penetrated across the rostral border of the lesion (Figures 2C”). In contrast, axons frequently penetrated the rostral border and extended rostrally in kaBRAF/PTEN iTg mice (Figure 2D”, Figure 2D’”). In addition, more numerous axons extended rostrally along the spinal cord surface in kaBRAF/PTEN iTg mice than in pre-conditioning mice (Figure 2C’”, Figure 2D’”). Regenerating DC axons preferentially extended rostrally along the spinal cord surface in both groups, but more frequently penetrated the CNS parenchyma in kaBRAF/PTEN iTg mice (Figure 2C’”, Figure 2D’”). Axon quantification revealed no significant difference in regenerating DC axon numbers within the lesion epicenter between kaBRAF/PTEN iTg and the pre-conditioning group (Figure 2E). Importantly, kaBRAF/PTEN iTg mice showed a significant increase in regenerating DC axons between 250 – 2000 μm across the lesion site compared to the pre-conditioning group. Quantification of regenerating DC axons within the CNS parenchyma also confirmed a significant increase between 250 – 2000 μm across the lesion epicenter in kaBRAF/PTEN iTg mice compared to the pre-conditioning group (Figure 2E).

### Supplemental pre-conditioning lesion enhances further the stimulatory effect of concomitant B-RAF activation and PTEN deletion

To determine whether there is an additive effect of concurrent B-RAF activation and PTEN deletion with pre-conditioning lesion, we crushed the right sciatic nerves at 10 days before SCI in another group of kaBRAF/PTEN iTg mice (kaBRAF/PTEN iTg + Pre-Conditioning, Figure 3A). The lesion area in kaBRAF/PTEN iTg + Pre-Conditioning mice ranged between 0.079 – 0.116 mm^2^ and was not significantly different compared to kaBRAF/PTEN iTg mice (Figure 3B). Notably, in contrast to mice with kaBRAF/PTEN iTg or pre-conditioning alone, all kaBRAF/PTEN iTg + Pre-Conditioning mice showed lesion epicenters that were densely populated by numerous DC axons that crossed the astroglial border caudal to the lesion site (Figure 3C’, Figure 3D’). Although most axons extended along the spinal cord surface, more axons penetrated the rostral astroglial border in kaBRAF/PTEN iTg + Pre-Conditioning mice than in the kaBRAF/PTEN iTg group (Figure 3C”, Figure 3D”). The number of regenerating DC axons extending rostrally along the spinal cord surface was also higher in the kaBRAF/PTEN iTg + Pre-Conditioning group than in kaBRAF/PTEN iTg mice and these axons often penetrated the CNS parenchyma (Figure 3C’”, Figure 3D’”). Axon quantification revealed a significant increase in DC axon number at 250 μm caudal to the lesion and within the lesion epicenter in the kaBRAF/PTEN iTg + Pre-Conditioning group compared to the kaBRAF/PTEN iTg group (Figure 3E). However, there was no statistically significant increase in the number of regenerating DC axons rostral to the lesion, perhaps because of the great variability among mice (Figure 3E). Similarly, quantification of DC axons within the CNS parenchyma showed no statistically significant difference between kaBRAF/PTEN iTg + Pre-Conditioning and kaBRAF/PTEN iTg groups (Figure 3E). We interpret these results to suggest that a pre-conditioning lesion has an additive role on the stimulatory effect of B-RAF activation and PTEN deletion, resulting in more extensive DC axon regrowth. However, most axons preferentially extended along the spinal cord, rather than directly penetrating the rostral border of the lesion.

**Figure 3.**
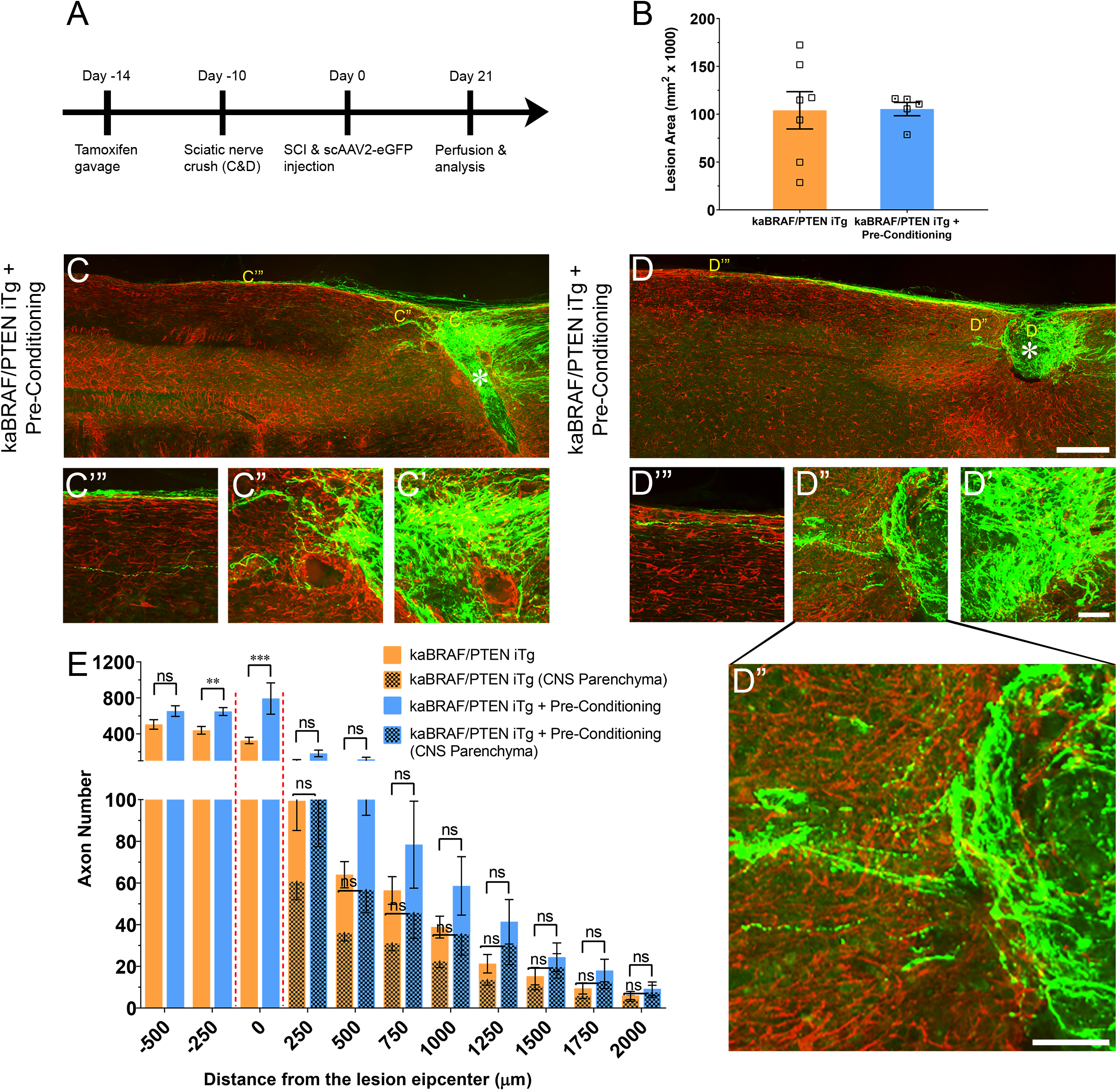
Additive effect of pre-conditioning lesion and concomitant B-RAF activation and PTEN deletion in promoting DC axon regeneration. (A) Schematic diagram showing the experimental timeline. Four days after the first tamoxifen gavage a group of kaBRAF/PTEN iTg mice underwent right sciatic nerve crush (kaBRAF/PTEN iTg + Pre-Conditioning). Both kaBRAF/PTEN iTg + Pre-Conditioning and kaBRAF/PTEN iTg mice then underwent dorsal SCI and scAAV2-eGFP injection into the right L4 and L5 DRGs to label regenerating DC axons two weeks after the first tamoxifen gavage. (B) Quantitative analysis of the lesion area after SCI in kaBRAF/PTEN iTg versus kaBRAF/PTEN iTg + Pre-Conditioning mice. (C–D’”) Representative images of two kaBRAF/PTEN iTg + Pre-Conditioning lesion mouse spinal cords showing a large number of regenerating DC axons immediately caudal to the lesion and numerous axons within the lesion epicenter. (E) Quantitative comparison of regenerated DC axons in kaBRAF/PTEN iTg group versus kaBRAF/PTEN iTg + Pre-Conditioning lesion group. Significantly increased DC axon regeneration is evident at 250μm caudal to the lesion in the combined group (**P=0.0037, df=10), and at the lesion epicenter (0μm: ***P<0.001, df=10). Two-way ANOVA with repeated measures. n= 7 (kaBRAF/PTEN iTg) and 5 (kaBRAF/PTEN iTg + Pre-Conditioning). (E, patterned) Quantitative comparison of regenerated DC axons within the CNS parenchyma rostral to the SCI in kaBRAF/PTEN iTg versus kaBRAF/PTEN iTg + Pre-Conditioning group. Scale bars, 250μm (C and D), 50μm (C’–C’”, and D’–D’”), ns, not significant.

### Co-targeting B-RAF and PTEN immediately after SCI promotes DC axon regeneration

To model a clinically relevant setting more closely, we next examined whether concomitant B-RAF activation and PTEN deletion immediately post-SCI can promote DC axon regeneration. To specifically activate B-RAF and delete PTEN in sensory neurons, we injected scAAV2-Cre unilaterally into the L4 and L5 DRGs in *LSL-kaBRAF:PTEN^f/f^:R26-TdT* transgenic mice that carry floxed alleles for kaBRAF with PTEN (hereafter as kaBRAF/PTEN/TdTom^+/+^). This transgenic line also expresses Rosa 26-TdTomato (*R26-TdT*) upon Cre-mediated recombination, which enables identification of fluorescent DC axons. We used *R26-TdT* (hereafter as TdTom^+/+^) mice as control. We first confirmed the efficiency of scAAV2-Cre-mediated recombination and transfection using TdTom^+/+^ mice (Supplementary Figure 2). Single unilateral microinjection of scAAV2-Cre into L4 and L5 DRGs resulted in high TdTom expression in transfected neuron somata (Supplementary Figure 2A) and DC axons (Figure 4C), confirming effective Cre-mediated recombination. To examine transfection efficiency, we co-stained microinjected DRGs with a mature neuronal marker (NeuN) and found that 63.6% (1351/2123, n = 3 mice, 6 DRGs) of sensory neurons were transfected (Supplementary Figure 2A, Supplementary Figure 2B). We further characterized scAAV2-Cre-labeled DRG neuron subtypes using immunohistochemistry (Supplementary Figure 2A, Supplementary Figure 2C). TdTom^+^ DRG neurons represented 80.4% (1132/1408) of large-diameter NF^+^ neurons, 39.6% (349/881) of CGRP^+^ peptidergic neurons, and 14.9% (126/848) of IB4^+^ non-peptidergic neurons (Supplementary Figure 2, Supplementary Figure 2C), suggesting that scAAV2-Cre predominantly transfects large-diameter sensory neurons. Single unilateral microinjection of scAAV2-Cre into kaBRAF/PTEN/TdTom^+/+^ mice DRGs also downregulated PTEN protein expression in sensory neurons, further confirming efficient Cre-mediated recombination (Supplementary Figure 2D).

**Figure 4.**
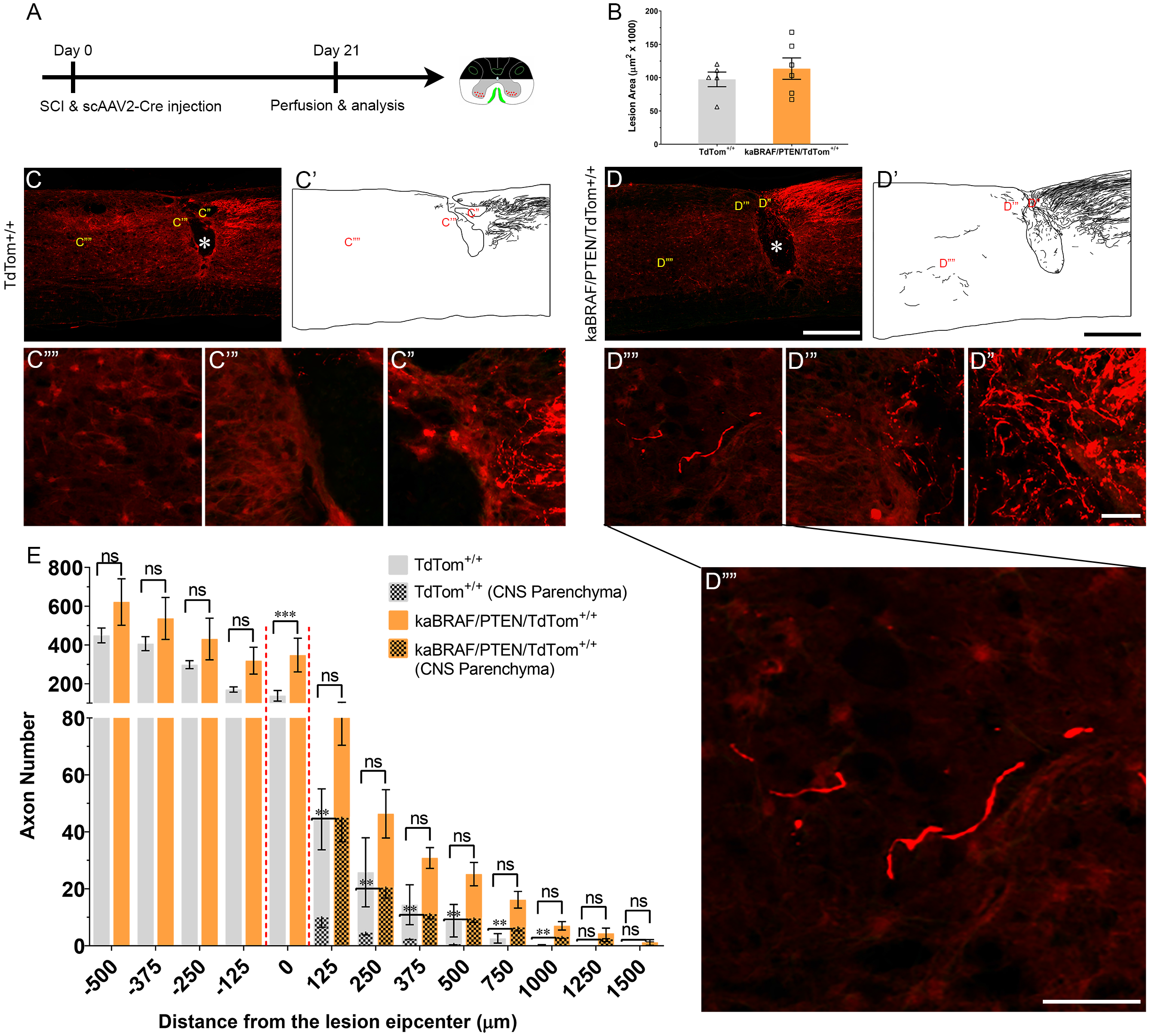
Co-targeting of B-RAF and PTEN immediately after SCI promotes DC axon regeneration. (A) Schematic diagram showing the experimental timeline. Immediately following dorsal SCI scAAV2-Cre was injected into the right L4 and L5 DRGs to induce B-RAF activation and PTEN deletion in *LSL-kaBRAF:PTEN^f/f^:R26-TdT* mice. scAAV2-Cre induction is also used to label regenerating DC axons via TdTomato expression upon Cre-mediated recombination in experimental (*LSL-kaBRAF:PTEN^f/f^:R26-TdT*) and control (*R26-TdT*) mice. (B) Quantitative analysis of the SCI lesion area in *LSL-kaBRAF:PTEN^f/f^:R26-TdT* versus *R26-TdT* mice. (C–C””) Representative images of a *R26-TdT* mouse spinal cord showing lack of DC axon regeneration within the lesion epicenter. Most axons stop at the caudal astroglial border of the injury site and rarely penetrate the lesion epicenter. (D–D””) Representative images of an *LSL-kaBRAF:PTEN^f/f^:R26-TdT* mouse spinal cord showing increased DC axon regeneration. Some regenerating DC axons penetrate the lesion epicenter, and a few extend directly across the rostral astroglial border of the injury site. (E) Quantitative comparison of regenerated DC axons in *LSL-kaBRAF:PTEN^f/f^:R26-TdT* versus *R26-TdT* mice. Significantly increased DC axon regeneration is evident at the lesion epicenter (0μm: ****P<0.0001, df=9). Two-way ANOVA with repeated measures. (E, patterned) Quantitative comparison of regenerated DC axons within CNS parenchyma rostral to the DC crush in *LSL-kaBRAF:PTEN^f/f^:R26-TdT* versus *R26-TdT* mice. Regenerating DC axons are significantly more numerous in *LSL-kaBRAF:PTEN^f/f^:R26-TdT* mice at distances >1000μm rostral to the lesion. 125μm: **P=0.0082, df=9; 250μm: **P=0.0079, df=9; 375μm: **P=0.0046, df=9; 500μm: **P=0.0019, df=9; 750μm: **P=0.0058, df=9; 1000μm: **P=0.0075, df=9; two-way ANOVA with repeated measures. n=5 (*R26-TdT*), and 6 (*LSL-kaBRAF:PTEN^f/f^:R26-TdT*). Scale bars, 500μm (C–C’, and D–D’), 50μm (C’–C””, and D”–D””), ns, not significant.

We then injected scAAV2-Cre into the L4 and L5 DRGs immediately after SCI in TdTom^+/+^ control and kaBRAF/PTEN/TdTom^+/+^ mice (Figure 4A). The lesion area ranged between 0.056 – 0.120 mm^2^ in TdTom^+/+^ control and 0.067 – 0.168 mm^2^ in kaBRAF/PTEN/TdTom^+/+^ mice (Figure 4B). Quantification of the lesion size showed no differences in lesion area between TdTom^+/+^ and kaBRAF/PTEN/TdTom^+/+^ groups (Figure 4B). At 3 weeks after SCI in TdTom^+/+^ mice, most DC axons stopped at the astroglial border caudal to the injury site. DC axons were rarely observed in the lesion epicenter (Figure 4C – C’). Most DC axons in TdTom^+/+^ animals displayed characteristic dieback from the lesion site and individual axons had several retraction bulbs (Figure 4C”). In all TdTom^+/+^ mice examined, DC axons failed to extend a notable distance rostrally across the astroglial border (Figure 4C’” – Figure 4C””). Remarkably, in kaBRAF/PTEN/TdTom^+/+^ animals many DC axons penetrated the lesion epicenter and several axons extended considerable distances rostrally over 1000 μm (Figure 4D – Figure 4D””). Individual DC axons in kaBRAF/PTEN/TdTom^+/+^ mice exhibited a distinctive regenerative morphology, including tortuous trajectories (Figure 4D””). Quantification of DC axons revealed a significant increase in regenerating DC axons at the lesion epicenter in kaBRAF/PTEN/TdTom^+/+^ mice compared to the TdTom^+/+^ group (Figure 4E). Rostral to the lesion, around 50% of regenerating DC axons in kaBRAF/PTEN/TdTom^+/+^ mice entered the CNS parenchyma (Figure 4D””). Quantification of regenerating DC axons in the CNS parenchyma showed a significant increase in axons that grew more than 1000 μm across the lesion epicenter in kaBRAF/PTEN/TdTom^+/+^ mice compared to the TdTom^+/+^ group (Figure 4E). Altogether, these results demonstrate that scAAV2-Cre predominantly labels large-diameter sensory neurons and that concomitant B-RAF and PTEN targeting immediately post-SCI enhances DC axon regeneration, although it is less vigorous compared to animals that were transduced 2 weeks before injury.

## DISCUSSION

Dorsal column crush causes complete destruction of the dorsal spinal cord parenchyma and totally disrupts ascending DC axons (Zhang et al., 1996). In line with earlier studies (Inman and Steward, 2003, Cafferty et al., 2004), we observed only a few DC axons approaching and entering the site of injury in WT mice (Figure 1C). In contrast, mice with concomitant B-RAF activation and PTEN deletion either before or immediately after SCI demonstrated many more axons regenerating into the lesion and densely populating its epicenter. Although most of these regenerating axons preferentially grew along the surface of the spinal cord, many extended directly across and beyond the lesion, and some even reached distances greater than 2000 μm rostral to the lesion by 3 weeks after SCI. These observations demonstrate that co-targeting B-RAF and PTEN powerfully enhances the intrinsic capacity of DC axons to regenerate after SCI.

Previous studies reported limited efficacy in promoting DC axon regeneration after SCI through targeting single (Seijffers et al., 2007, Bareyre et al., 2011, Yu et al., 2011, Wang et al., 2015, Wang et al., 2017) or multiple (Bomze et al., 2001, Fagoe et al., 2015) intrinsic pro-regenerative pathways. For instance, constitutive overexpression of activating transcription factor 3 (*Atf3*) failed to promote DC axon regeneration 4 weeks following thoracic 10 (T10) dorsal hemisection in mouse (Seijffers et al., 2007). Similarly, pre-injury overexpression of signal transducer and activator of transcription 3 (*Stat3*) increased DC axon sprouting at 2 days post-lesion but did not enhance their regeneration at 10 days after cervical dorsal hemisection in mice (Bareyre et al., 2011). Pre-injury overexpression of *Klf7* (Wang et al., 2017) or *Sox11* (Wang et al., 2015) reduced DC axonal dieback without inducing regenerative growth up to 8 weeks after cervical 5 (C5) dorsal hemisection in mice. Comparable results have been reported for rat. In one recent example, pre-injury overexpression of the proto-oncogene *Myc* prevented DC axon retraction but failed to promote their rostral outgrowth by 4 weeks after T9 dorsal hemisection (Shin et al., 2021). Equally, pre-injury overexpression of DNA binding 2 (*Id2*) enabled DC axons to reach the injury site without extending beyond the lesion epicenter by 2 weeks following T10 dorsal hemisection in mice (Yu et al., 2011). Constitutive neuronal overexpression of two growth cone proteins (GAP-43 and CAP-23) triggered DC axon regeneration over 5 mm into the sciatic nerve grafted within the lesion site by 4 months following cervico-thoracic dorsal hemisection in mice (Bomze et al., 2001). Notably, neither GAP-43 nor CAP-23 overexpression alone could promote DC axon regeneration into a graft (Bomze et al., 2001). However, simultaneous pre-injury overexpression of four transcription factors (*Atf3*, *c-Jun*, *Stat3*, and *Smad1*) failed to enhance DC axon regeneration 4 weeks after C4 dorsal hemisection in rats (Fagoe et al., 2015).

Pre-injury overexpression of CREB promoted DC axon regeneration into the lesion site by 4 weeks after T6-T7 dorsal hemisection in rats (Gao et al., 2004). Similarly, overexpression of BMP4 either before injury or immediately post-SCI triggered DC axon regeneration into the lesion epicenter but not rostral to the lesion by 4 weeks after T8 dorsal hemisection in mice (Parikh et al., 2011). Equally, enforced α9 integrin expression immediately post-SCI stimulated DC axon regeneration into the lesion epicenter by 6 weeks following C5/C6 dorsal column crush in rats (Andrews et al., 2009).

Among the multiple axon growth repressors identified to date, only GSK3β has been targeted to enhance DC axon regeneration after SCI. Constitutive complete or partial GSK3β deletion promoted DC regeneration that extended within the lesion site by 5 weeks after T9/T10 dorsal hemisection in mice (Liz et al., 2014). Co-deleting two intrinsic axon growth repressors, SOCS3 and PTEN, induced modest peripheral sensory axon regeneration after sciatic nerve crush in mice (Gallaher and Steward, 2018); whether this strategy will enhance DC axon regeneration after SCI is unknown.

Other studies targeting extrinsic inhibitors (Lee et al., 2010, Shields et al., 2008) and intrinsic and extrinsic molecules in combination (Wang et al., 2017) also reported promoting DC axon regeneration after SCI with moderate efficacy. Sustained chondroitinase ABC (ChABC) release within the lesion site blocked chondroitin sulfate proteoglycans (CSPGs)-mediated inhibition and prevented DC axonal dieback but failed to promote their rostral re-growth at 6 weeks after T10 dorsal hemisection in rats (Lee et al., 2010). Intrathecal ChABC injection mildly enhanced DC axon regeneration towards the lesion site, but only a few axons extended rostrally at 5 weeks after C3 dorsal hemisection in rats (Shields et al., 2008). Combined intraparenchymal ChABC injection and *Klf7* overexpression stimulated DC axon regeneration towards the lesion site without enhancing their regenerative growth within or across the lesion site by 8 weeks after C4/C5 dorsal hemisection in mice (Wang et al., 2017). Concomitant ChABC and neurotrophin-3 treatments induced DC axon extension beyond 500 μm rostral to the lesion at 6 weeks following T10 dorsal over-hemisection in rats (Lee et al., 2010). Our observation that DC axons regenerate farther than 2000 μm rostral to a bilateral dorsal column crush injury by 3 weeks highlights the increased potency of concomitantly targeting B-RAF and PTEN, although differences in injury models may also contribute to these differences.

The mechanisms by which B-RAF activation with PTEN deletion powerfully and synergistically promotes DC axon regeneration remain unclear. A recent transcriptomic study combining data from several independent laboratories has provided a comprehensive list of regeneration-associated genes (RAGs) that are responsible for sensory axon regeneration after pre-conditioning lesion (Chandran et al., 2016). *In silico* examination of this list did not implicate a downstream mediator of B-RAF as a RAG, including *Braf*, *Mapk1*, *Map3k2*, *Map2k1*, *Creb1*, and *Rps6ka1* (Chandran et al., 2016). Downstream mediators of PTEN, including *Rps6*, *Rps6kb1*, and *Mtor* were also not identified as RAGs, suggesting that pre-conditioning lesion and B-RAF/PTEN co-targeting depend on distinct underlying mechanisms. In addition, as mentioned earlier, enforced expression of multiple identified RAGs (*Sox11*, *Atf3*, *Jun*, *Stat3*, and *Smad1*) in sensory neurons either individually (Seijffers et al., 2007, Bareyre et al., 2011, Wang et al., 2015) or in combination (Fagoe et al., 2015) did not promote robust DC axon regeneration after SCI, suggesting that different molecules are responsible. Furthermore, our findings that concomitant B-RAF and PTEN targeting induces significantly greater DC axon regeneration than a pre-conditioning lesion (Figure 2), and that a supplemental pre-conditioning lesion has an additive, rather than a synergic, effect on B-RAF/PTEN-mediated DC axon regeneration (Figure 3), provide additional evidence that their underlying mechanisms are likely to differ. Consistent with this notion, pre-conditioning injury in kaBRAF/PTEN iTg animals increased DC axon regeneration in the same region as in WT mice: within and 250 μm caudal to the lesion. Therefore, it remains an important challenge to identify growth-promoting pathways or downstream signaling effectors that are uniquely activated upon concomitant B-RAF activation and PTEN deletion in the injured sensory neurons. These studies may reveal novel signaling pathways or effectors that drive the remarkable synergistic effect of concomitant B-RAF and PTEN targeting on DC axon regeneration.

## MATERIALS AND METHODS

### Animals

All procedures, including animal care and maintenance, complied with the National Institute of Health guidelines regarding the care and use of experimental animals and were approved by the Institutional Animal Care and Use Committee of Temple University, Philadelphia, PA, USA. Male and female mice, aged 6 weeks and weighing over 20 g, were used in all experiments and were maintained on the C57/BL6 background. *LSL-kaBRAF:PTEN^f/f^:brn3a-CreER^T2^:R26-TdT* (kaBRAF/PTEN iTg) transgenic lines were kindly provided by Jian Zhong (Cornell University). The *LSL-kaBRAF* and *Brn3a-CreER^T2^* deleter mouse lines were generated and genotyped as described previously (O’Donovan et al., 2014, Eng et al., 2001). Cre-negative (*LSL-kaBRAF:PTEN*) mice from the same litter were used as control (WT). Transgenic *Rosa26(R26)-TdT* mice were purchased from the Jackson Laboratory (Stock # 007914).

### Genotyping

Genotyping was performed by polymerase chain reaction (PCR) using the following primer pairs: *Braf* = 5’-GCCCAGGCTCTTTATGAGAA-3’ (common forward), 5’-AGTCAATCATCC ACAGAGACCT-3’ (reverse, mutant allele), 5’GCTTGGCTGGACGTAAACTC-3’ (reverse, wildtype allele), *Rosa26* = 5’-AAGGGAGCTGCAGTGGAGTA-3’ (forward, wildtype allele), 5’-CCGAAAATCTGTGGGAAGTC-3’ (reverse, wildtype), 5’-GGCATTAAAGCAGCATATCC-3’ (reverse, mutant allele), 5’-CTGTTCCTGTACGGCATGG-3 (forward, mutant allele), *Brn3aCre* = 5’-CGCGGACTTTGCGAGTGTTTTGTGGA-3’ (forward), 5’-GTGAAACAGCATTGCTGTCACTT-3’ (reverse), *Pten* = 5’-CAAGCACTCTGCGAACTGAG-3’ (forward), and 5’-AAGTTTTTGAAGGCAAGATGC-3’ (reverse).

### Tamoxifen administration

A 10 mg/ml tamoxifen solution was prepared in 10:1 sunflower oil:100% ethanol. 50 μl tamoxifen (10 mg/ml) was given via oral gavage for 5 consecutive days, 2 weeks prior to injury. To account for the possible effects of tamoxifen, we also applied the same tamoxifen administration paradigm to WT and pre-conditioning groups.

### Surgical procedure

Two weeks following the first tamoxifen gavage, mice underwent dorsal column injury. They were anesthetized by intraperitoneal (i.p.) injection of ketamine (8 mg/kg), and xylazine (120 mg/kg). Supplements were given during the surgical procedure as needed. Small animal hair clippers (Oster Professional Products) were used to remove the hair overlying the surgical site. A 2-3 cm incision was made in the skin overlying the thoracic region, the spinal musculature was reflected, and the T10-T11 spinal cord segments were exposed by laminectomy. The cavity made by the laminectomy was continuously perfused with warm sterile Ringer’s solution. Lidocaine (2%) was added dropwise (~2-3 drops) to the exposed spinal cord region. The dorsal part of the spinal cord was completely crushed for 2 s with a modified #5 forceps (Dumont, Fine Science Tools, Inc., Foster City, CA, USA). The 4–5 mm of the forceps adjacent to the tip were thinned to a width of 0.1 mm. The spinal dura and the dorsal spinal vein remained intact after crushing. For pre-conditioning lesion, the right sciatic nerve was crushed 10 days prior to the SCI, as we described previously (Di Maio et al., 2011). Following SCI, a piece of biosynthetic membrane (Biobrane, UDL Laboratories, Inc., Sugarland, TX, USA) was placed over the injured site. Muscles were then sutured with sterile 5-0 sutures (Ethicon, Cincinnati, OH, USA), and the midline incision closed with wound clips (Fine Science Tools, Inc., Foster City, CA, USA).

### Adeno-associated virus vector injection

Recombinant self-complementary adeno-associated virus serotype 2 expressing enhanced green fluorescent protein under the control of a CMV-enhanced chicken β-actin (CAG) promoter (scAAV2-eGFP, Vector Biolabs, catalogue #7072, 1×10^13^ genomic copy/ml) was used to label regenerating DC axons. We also used scAAV2 carrying Cre (scAAV2-Cre, 1×10^12^ genomic copy/ml kindly provided by George M. Smith, Temple University) to transduce DRG neurons in *LSL-kaBRAF:PTEN^f/f^:R26-TdT* and *R26-TdT* mice. Immediately after SCI, the right L4 and L5 DRGs were exposed, and the mice were placed in a stereotaxic holder with spinal cord clamps (STS-A, Narishige Group, Tokyo, Japan). Either scAAV2-eGFP or scAAV2-Cre was microinjected using a micropipette pulled to a diameter of 0.05 mm and a nanoinjector (World Precision Instruments, Inc., Sarasota, FL, USA). For each injection, the micropipette was introduced 0.5 mm into the DRGs, and 1 μl of the virus was injected over a 10 min period (100 nl/min). The glass needle was left in place for 3 additional min following each injection to allow adequate virus diffusion within the DRGs. Following virus injection, a piece of biosynthetic membrane (Biobrane, UDL Laboratories, Inc., Sugarland, TX, USA) was placed over the exposed spinal cord and dura to minimize scar formation. The musculature was closed with 5-0 sterile silk sutures (Ethicon) and the skin with wound clips (Fine Science Tools, Inc., Foster City, CA, USA). To avoid dehydration, mice were given subcutaneous injections of lactated Ringer’s solution and kept on a heating pad until fully recovered from anesthesia. Buprenorphine (0.05 mg/kg) was given intramuscularly for 2 consecutive days after SCI.

### Immunohistochemistry, fluorescence, and confocal microscopy

Three weeks following SCI and virus injection, the mice were anesthetized with a lethal dose of 10% Euthasol (Virbac, Westlake, TX, USA) in 0.9% saline and perfused transcardially with 4% paraformaldehyde (PFA) in PBS. The spinal cord containing injected DRs was dissected out, post-fixed in 4% PFA for 2 additional hrs and stored in 30% sucrose in 1× PBS, pH 7.4, overnight at 4°C for cryoprotection. Tissues were frozen-embedded in cryoprotectant medium (M-1 Embedding Matrix, Thermo Fisher Scientific, Waltham, MA, USA) in isomethylbutane at −80°C. Serial sagittal spinal cord sections (20 μl) were cut using a cryostat (Leica Microsystems, Wetzlar, Germany) and collected directly onto Superfrost Plus glass slides. To ensure the absence of spared DC axons, we examined all spinal cord and sagittal sections and confirmed that there were no GFP^+^ or TdTom^+^ axons over 4 mm rostral to the lesion site (data not shown). For immunolabeling, sections were rehydrated by rinsing in three changes, 10 min each, of 1× PBS, incubated in 0.1 M glycine in 1× PBS for 15 min, and subsequently blocked in 0.2% Triton X-100, 2% bovine serum albumin (BSA) in 1× PBS for 15 min. Sections were incubated with primary antibodies overnight at 4°C, washed with 2% BSA in PBS three times, 10 min each, and incubated with secondary antibodies for 1 hr. Primary antibodies were used at the following concentrations for immunohistochemistry: rabbit anti-CGRP (1:1000, Peninsula Labs, Burlingame, CA, USA, T-4032), mouse anti SMI312 (1:1000, Biolegend, San Diego, CA, USA, 83790), chicken anti-GFP (1:1000, Avés Labs Inc., Tigard, OR, USA, 1020), rabbit anti-GFAP (1:500, Dako, Santa Clara, CA, USA, Z0334), mouse anti-GFAP (1:500, Sigma-Aldrich, Burlington, MA, USA, G3893), mouse anti NeuN (1:1000, MilliporeSigma, Burlington, MA, USA, MAB377), rabbit anti red fluorescent protein (1:5000, Rockland, Limerick, PA, USA, 600-401-379), and rabbit anti-PTEN (1:200, Cell Signaling, Danvers, MA, USA, 9559). Secondary antibodies used were Alexa Fluor 647 goat anti-mouse IgG1 (1:400, Molecular Probes, Eugene, OR, USA, A-21240), Alexa Fluor 568 goat anti-rabbit IgG (1:400, Molecular Probes, Eugene, OR, USA, A-11011), Alexa Fluor 647 goat anti-rabbit IgG (1:400, Molecular Probes, Eugene, OR, USA, A-21244), Fluorescein-conjugated donkey anti-rabbit IgG (1:400, Jackson ImmunoResearch Labs Inc., West Grove, PA, USA, 711-096-152), Fluorescein-conjugated goat anti-rabbit IgG (1:400, Chemicon International, Temecula, CA, USA, AP307F), and Alexa Fluor 488 donkey anti-chicken IgG (1:400, Jackson ImmunoResearch Labs Inc., West Grove, PA, USA, 703-545-155).

For Isolectin B4 (IB4) immunostaining, sections were first post-fixed in 4% PFA, rehydrated in 1× PBS, quenched in 0.1 M glycine in 1× PBS, and permeabilized with 0.2% Triton X-100/PBS (PBST). Sections were then incubated with an IB4-biotin conjugate (1:200, Sigma-Aldrich, Burlington, MA, USA, L2140) prepared in 5% normal goat serum (NGS)/PBST overnight at room temperature. Sections were then blocked with 5% NGS/PBS and incubated with rhodamine (TRITC) streptavidin (1:400, Jackson ImmunoResearch Labs Inc., West Grove, PA, USA, 016-020-084) prepared in 1× PBS for 1 hr. DAPI nucleic acid stain (1:1000, Invitrogen, Waltham, MA, USA, D-1306) was used to counterstain prior to the final washes in 1× PBS. All antibodies were diluted in 2% BSA in PBS. Sections were washed in three changes of 1× PBS, 10 min each, and mounted with Vectashield mounting medium (Vector Laboratories, Burlingame, CA, USA).

Conventional fluorescence microscopy was performed using Olympus BX53 and Zeiss Axio Imager microscopes. Images were captured using a Zeiss Axio Imager upright fluorescence microscope with a 10 × 0.45 NA or 20 × 0.8 NA objective. Z stacked images were acquired using the AxioVision (Zeiss) software. We also used a SP8 confocal microscope (Leica Microsystems, Wetzlar, Germany) with either a 40 × 1.3 NA or 63 × 1.4 NA objective and Leica proprietary software. Acquired stacks were assembled using the maximum projection tool. All images were processed using Imaris (Bitplane, Zürich, Switzerland), and figures were prepared using Adobe Photoshop (Adobe, San Jose, CA, USA).

### Quantification of scAAV2-Cre labeled neurons

The number of NeuN/scAAV2-Cre, SMI312/scAAV2-Cre, CGRP/scAAV2-Cre, and IB4/scAAV2-Cre co-labeled neurons in L4 and L5 DRGs was counted in 20 non-adjacent sections from 3 independent mice. Co-labeled neurons were averaged from all 20 sections for each mouse.

### Quantification of regenerating DC axons and lesion area after SCI

We used a consecutive series of 20 μm thick sagittal sections to determine the total number of regenerating DC axons at different distances from the lesion epicenter, which was identified based on GFAP immunostaining (Figure 1C). Perpendicular lines were drawn on to the sagittal plane at the lesion epicenter and at 0.25 mm intervals from 0.5 mm caudal to 2 mm rostral to the lesion epicenter. All GFP^+^ or TdTom^+^ axons crossing the perpendicular lines at each distance were counted manually using the ImageJ Cell Counter plug-in. The total number of axons at a given distance was summed from all immunostained sections containing GFP^+^ or TdTom^+^ axons. Based on overlay images of GFP and GFAP or TdTom and GFAP, regenerating axons in the rostral segments of the spinal cord were sub-divided into two groups: (i) associated with GFAP-positive astrocytes: either grew directly across the lesion site or entered the GFAP-positive territories of the spinal cord above the lesion site, and (ii) not associated with GFAP-positive astrocytes: circumvented the lesion site and grew along the surface of the spinal cord. Lesion area was quantified using ImageJ based on GFAP staining in all consecutive sagittal sections in which regenerating DC axons were quantified. Lesion area was quantified based on GFAP staining using ImageJ software on 7-10 sagittal sections and then averaged to estimate the final lesion area in μm^2^.

### Statistical analysis

Statistical significance of axon regeneration was assessed by 2-way ANOVA with repeated measures using Statistical Analysis System (SAS) software (SAS Institute Inc., Cary, NC, USA) and GraphPad Prism 6.0 (GraphPad, San Diego, CA, USA). Data were split between rostral and caudal to the lesion before carrying out 2-way ANOVA with repeated measures. Lesion area was assessed by un-paired t-test using GraphPad Prism 6.0 (GraphPad, San Diego, CA, USA). All data are presented as mean ± standard error of the mean (SEM). Sample sizes are as described in the figure legends. Results were considered statistically significant if the p-value was < 0.05.

## COFLICT OF INTEREST

The authors declare that the research was conducted in the absence of any commercial or financial relationships that could be construed as a potential conflict of interest.

## AUTHOR CONTRIBUTIONS

HNN: Data curation, Formal analysis, Writing - original draft

HK: Investigation, Methodology

SP: Resources, Investigation

JZ: Resources, Methodology

YS: Conceptualization, Supervision, Funding acquisition, Investigation, Project administration, Writing - review and editing

## FUNDINGS

This work was supported by Shriners Children’s (257487 and 268980 to Y-J.S.; 84600 to H.N.N.), and NIH NINDS (NS253020 to Y-J.S.).

## ACKNOWLEDGMENTS

We thank all members of Son laboratory for critical reading of the manuscript. We also thank Dr. Huaqing Zhao, and Xiaoning Lu for statistical analysis.

---

**Supplementary Figure 1. Serial sections from the kaBRAF/PTEN iTg mouse spinal cord shown in Figure 1D**

Numerous regenerating sensory axons penetrate the lesion epicenter and extend long distances rostral to the lesion site at 3 weeks after SCI. DC axons are labeled by injecting scAAV2-eGFP into right L4 and L5 DRGs. #16 image is also shown in Figure 1D. Scale bar: 200μm.

**Supplementary Figure 2. Viral and genetic targeting of B-RAF and PTEN**

(A) Representative image of a *R26-TdT* mouse DRG microinjected with scAAV2-Cre and co-stained with the global mature neuronal marker NeuN, large-diameter neuron marker NF, small-diameter peptidergic neuron marker CGRP, and small diameter non-peptidergic neuron marker IB4. (B) Quantitative analysis of transfected (NeuN^+^/TdTom^+^, arrowheads in A) mature neurons as a percentage of total (NeuN^+^) DRG neurons. Single unilateral microinjection of scAAV2-Cre into *R26-TdT* mice transfected >63.6% of mature neurons in the DRG. (C) Quantitative analysis of transfected large diameter (NF^+^/TdTom^+^), peptidergic (CGRP^+^/TdTom^+^), and non-peptidergic (IB4^+^/TdTom^+^, arrowheads in A) DRG neurons. Single unilateral microinjection of a scAAV2-Cre into *R26-TdT* mice transfected >79.4% of NF^+^ large-diameter neurons, 39.6% of small-diameter peptidergic CGRP^+^ neurons, and 15% of small-diameter non-peptidergic IB4^+^ neurons in the DRG. (D) Representative image of an *LSL-kaBRAF:PTEN^f/f^:R26-TdT* mouse DRG microinjected with scAAV2-Cre and co-stained with the PTEN antibody showing high TdTom expression and PTEN deletion (arrowheads) in sensory neurons upon Cre-mediated recombination. PTEN deletion is particularly evident in large-diameter neurons. Scale bar, 50μm (A).

## Notes

### Competing Interest Statement

The authors have declared no competing interest.

