## Supplementary figures and images for "Co-targeting B-RAF and PTEN enables sensory axons to regenerate across and beyond the spinal cord injury"

### Supplementary Figure 1 and Supplementary Figure 2

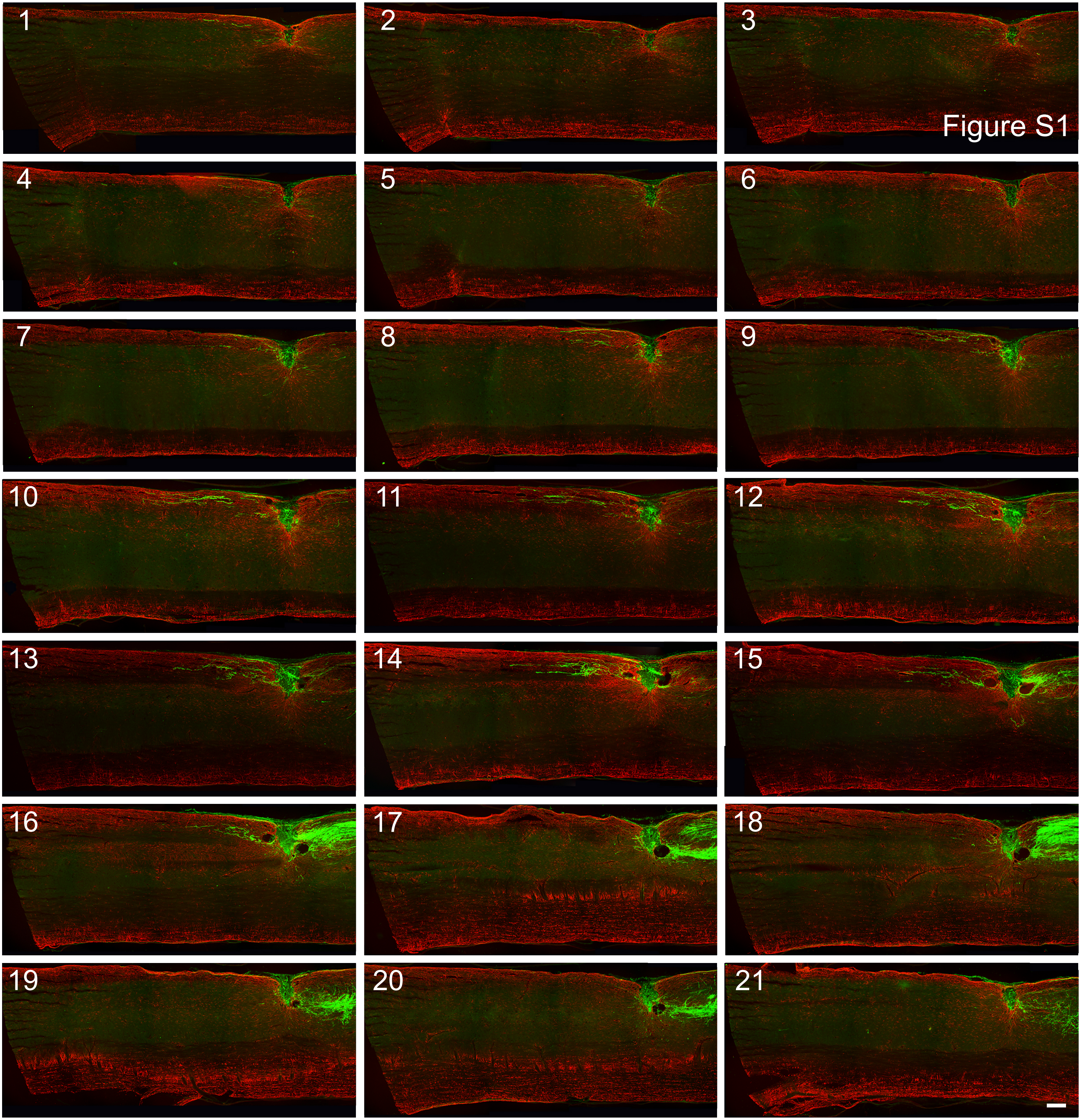

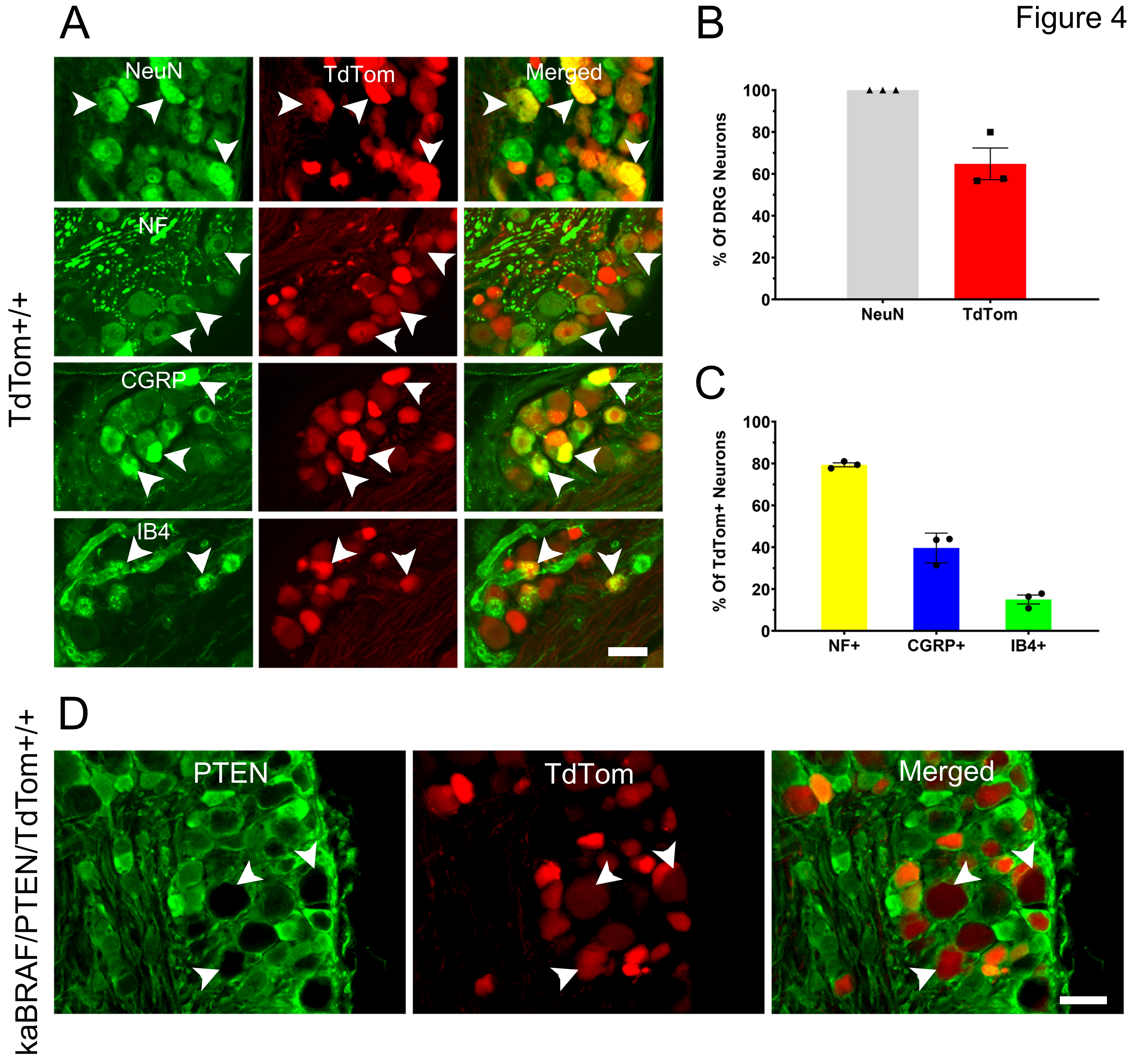
